# qcMol: a large-scale dataset of 1.2 million molecules with high-quality quantum chemical annotations for molecular representation learning

**DOI:** 10.1101/2025.09.07.674462

**Authors:** Haoyu Wang, Ziyan Zhang, Haipeng Gong

## Abstract

Recent advancements in deep learning have greatly prompted the *de novo* design of drugs and materials. Previous studies have shown that a well-designed molecular representation is critical for improving the accuracy of deep-learning-based molecular property prediction methods. However, the lack of large-scale data enriched with detailed physicochemical information hinders effective learning of an informative molecular representation. To fill this data gap, we introduce qcMol, a dataset consisting of 1.2 million molecules from 95 datasets with high-quality quantum chemical annotations, to facilitate molecular representation learning as well as downstream molecular property prediction. Chemicals in this dataset include drug-like compounds, metabolites and molecules with matched experimental data, covering 247,448 kinds of scaffolds and a broad spectrum of molecular sizes. Each compound in qcMol is annotated with detailed quantum chemical information, obtained through reliable quantum chemical calculations based on B3LYP-D3/def2-SV(P)//GFN2-xTB as well as the follow-up wave function post-analysis. These features are organized into multiple formats, allowing for flexible integration into diversified molecular representation learning frameworks. The broad data distribution, comprehensive quantum chemical annotations and flexible data formats jointly enable qcMol to serve as the pre-training resource as well as the benchmark test set for deep learning models, benefiting the practical *in silico* drug discovery.

qcMol is freely accessible from https://structpred.life.tsinghua.edu.cn/qcmol/.

## Introduction

The discovery of new drugs and materials is time-consuming and financially demanding. On average, this process takes 12 years from start to commercialization, costing 9 billion dollars^1–3^. Hence, previous developments in this field have been largely supported by the collaboration between physics-based calculations and laboratory experiments^4–6^. Benefiting from advances in artificial intelligence, various deep-learning-based methods have been developed, aiming to accelerate this process^7–10^. Compared with physics-based methods^11,12^, deep learning methods are advantageous not only in the improved performance but also in the markedly reduced inference time^13^. However, as a data-driven approach, the power of deep learning models is highly dependent on the quality of input data. Particularly, an informative and comprehensive molecular representation can improve the performance of deep learning models, effectively accelerating molecular discovery^14,15^. This principle has motivated the extensive investigation on molecular representation learning^10,16^.

Conventional molecular descriptors can be used as input features for representation learning models in multiple formats, including strings^17,18^, images^19^ and graphs^9^. Generated using cheminformatics tools like RDKit^20^ and Open Babel^21^, these descriptors are often too simplistic to capture the complex electronic structures of molecules, and conversely, the lack of physical accuracy in the representation of learning targets tends to misguide model training, potentially disrupting prediction power and generalizability^22,23^. Density functional theory (DFT)^24^, a reliable and computationally efficient quantum chemical method, provides access to electronic structures and relevant properties like electron density, molecular orbitals and energy levels. Wave functions obtained from such quantum chemical calculations can be localized using methods like Natural Bond Orbital (NBO)^25^ analysis to extract chemically interpretable electronic features, enabling the generation of structured representations that are inherently compatible with graph neural networks (GNN) and related frameworks without losing the quantum chemical fidelity. Moreover, incorporation of quantum chemical information, whether as explicit input features or through integrated training strategies, has been reported to benefit molecular property prediction tasks by the consideration of molecular electronic structures^26^. Hence, a well-maintained dataset composed of versatile molecular features derived through reliable quantum chemical calculations as well as the follow-up wave function post-analysis is essential for the training and benchmarking of deep-learning-based molecular representation learning and property prediction models.

In the past years, several quantum chemistry databases and platforms have been developed to support machine learning applications in molecular science^27–33^. Among them, QM9^32^ is one of the most widely used, containing computed electronic, energetic, geometric and thermodynamic properties for 134,000 stable small organic molecules that are composed of C, H, O, N and F elements and are chosen from GDB-17^34^, a dataset constructed with the purpose of exploring the chemical space. However, this dataset restricts the molecular size to be no more than nine heavy atoms, which substantially impairs the chemical diversity, and consequently is mainly used as a benchmark set for the evaluation of molecular property prediction models. Another relevant dataset, the PubChemQC Project^29^, provides ground-state electronic structures for over 3 million molecules computed by DFT at the B3LYP/6-31G^*^ level, as well as low-lying excited states of more than 2 million molecules calculated by time-dependent DFT (TD-DFT) with B3LYP/6-31+G^*^. Unfortunately, the involved molecules are randomly selected from the PubChem database, without prioritizing the relevance to drug discovery or material science applications. More importantly, this dataset primarily provides global molecular properties, such as the HOMO–LUMO gap and dipole moment, while lacking fine-grained electronic features at the levels of atoms or bonds. QuanDB^31^, a recently released dataset, incorporates 154,610 compounds composed of nine elements (H, C, O, N, P, S, F, Cl and Br) from public databases and literature. In addition to global molecular properties, it provides five local quantum chemical descriptors and the most stable 3D conformation for each molecule. However, the dataset covers a limited range of molecular sizes and features, lacking molecules of over 40 atoms and essential features.

In this work, to address the limitations of existing data resources, we constructed a new dataset named qcMol, which contains high-quality and abundant molecular features with distributions closely matching the real-world compounds, aiming to facilitate representation learning on chemical molecules. Unlike contemporary datasets that focus on small-molecule compounds, qcMol additionally includes drug-like molecules and common metabolites, many of which are associated with paired experimental measurements. Moreover, it provides detailed quantum chemical descriptors in a suitable format to simplify feature processing by graph-based deep learning frameworks. Incorporation of experimental results also allows qcMol to serve as a benchmark set for the training and refining of deep learning models developed for practical molecular design and drug discovery tasks.

## Methods

The qcMol computational workflow integrates several advanced computational chemistry tools, including RDKit^20^ (2022.09.5 version), Open Babel 3.1.1^21^, GFN2-xTB 6.5.1^35^, ORCA 5.0.3^36–41^, NBO 7.0^25^, and Multiwfn 3.8^42^. The complete dataset construction pipeline is described below and illustrated in Figure 1.

**Figure 1.**
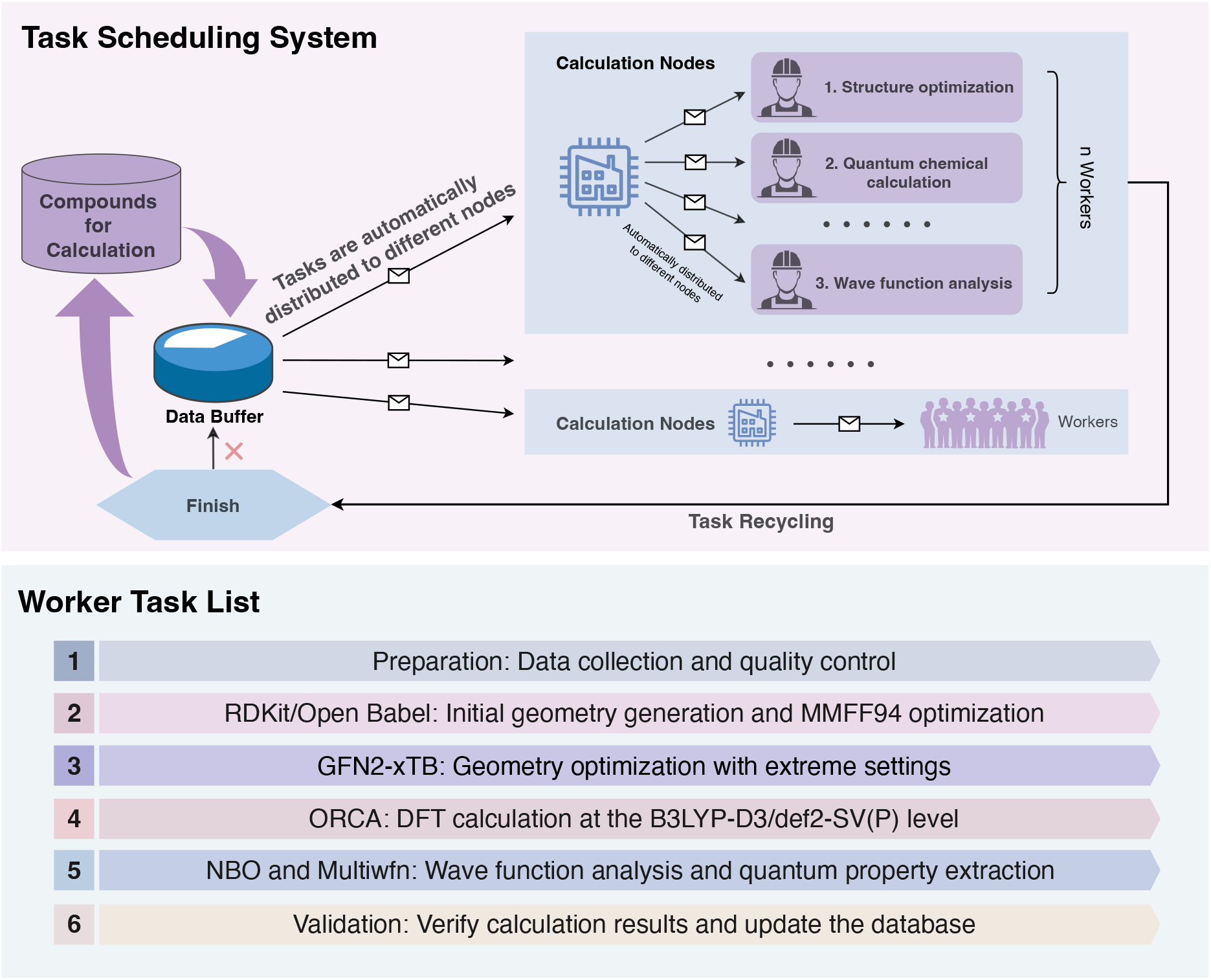
Task scheduling system and quantum chemical calculation pipeline of qcMol. Tasks are automatically distributed to computing nodes to enable parallel quantum chemical calculations for large-scale molecules.

### Dataset collection and curation

To fully utilize computational resources, we carefully selected the molecules for calculation and applied quality control procedures to the input data.

#### Data collection

qcMol integrates small-molecule data from various sources, including published literature and established biochemical databases. HMDB^43^, MoleculeNet^44^, MoleculeACE^45^, PDBbind^46^, and several other publicly available datasets^47^ are among the commonly used sources incorporated into qcMol. We collect each molecule’s identifier and associated experimental data while retaining as much relevant metadata as possible for downstream applications.

#### Data curation

Due to the limited computational resources and potential inconsistencies among the raw datasets, quality control procedures are necessary to ensure data reliability and usability. As molecular structures were recorded using SMILES, a data cleaning process was first applied to address inconsistencies in encoding formats and the lack of standardization (Figure 1). Specifically, we applied the following data cleaning criteria: 1) entries with inconsistent results assigned to identical SMILES strings were removed; 2) multi-component SMILES (*e*.*g*., representing mixtures or salts) were flagged for further processing; 3) molecules containing elements beyond the first four periods of the Periodic Table were filtered out; 4) entries containing metal ions were excluded; 5) all SMILES strings were converted into the RDKit-canonicalized SMILES, InChI, and InChIKey formats; 6) each molecule was assigned a unique internal identifier (*i*.*e*. qcMol ID) for reference within the database; 7) a deduplication step was conducted against existing entries, preventing redundant quantum chemical calculations. In qcMol, different conformers of the same molecule are treated as the same molecular entity, whereas molecules with different charge states are considered as distinct entries. Finally, to ensure the completeness of the database, we retrieved additional molecular metadata from the PubChem^48^ database for each compound.

### Workflow for quantum chemical calculations

Following standard notation in quantum chemistry, our computational scheme is denoted as B3LYP-D3/def2-SV(P)//GFN2-xTB, which indicates that single-point energy calculations using the B3LYP-D3^49,50^ functional with the def2-SV(P)^41^ basis set were performed on geometries optimized at the GFN2-xTB^35^ level (Figure 1).

#### Initial Structure Generation

Initial molecular structures were generated using RDKit^20^, an open-source cheminformatics toolkit. Specifically, RDKit was used to construct molecular geometries and perform preliminary optimization using the MMFF94 force field^51^. The resulting structures were saved in the XYZ format. If RDKit failed to generate a valid structure, the molecule was processed using Open Babel^21^ as a fallback. The generated structure served as the initial input for subsequent geometry optimization and quantum chemical calculations.

#### Preliminary Structure Optimization

In consideration of the trade-off between computational accuracy and efficiency, the initial structures were first optimized using GFN2-xTB before being passed to ORCA for higher-level quantum chemical calculations. GFN2-xTB is an extended tight-binding method parameterized from DFT, allowing efficient geometry optimization for large molecular systems composed of main-group elements. Specifically, the ANCopt algorithm was employed for geometry optimization, which iteratively minimized energy through structural refinement^35^. Notably, we adopted the “extreme” optimization level in xTB to maximize geometric precision while maintaining computational efficiency, yielding high-quality initial structures for subsequent quantum chemical calculations.

#### DFT-Based Quantum Chemical Calculations

Based on the optimized structures, advanced *ab initio* calculations were performed using ORCA^36–41^ at the B3LYP-D3 level with the def2-SV(P) basis set. B3LYP-D3 is a hybrid functional that combines Becke’s three-parameter exchange functional with the Lee-Yang-Parr correlation functional, and includes semi-empirical D3 dispersion corrections with atom-pair-specific parameters to account for long-range interactions accurately. The def2-SV(P) basis set is a double-*ζ* quality basis set that includes polarization functions.

#### Wave function Post-analysis

NBO 7.0^25^ and Multiwfn^42^ were employed to analyze the wave function obtained from ORCA comprehensively. The NBO method decomposes wave functions into localized bonding and non-bonding orbitals, providing a chemically meaningful picture of molecular structure and electron population. In addition, Multiwfn was used to compute a range of electronic properties, including electron density, molecular orbitals, partial atomic charges and bond-level descriptors.

The hierarchical strategy described above combines the computational efficiency of xTB-based geometry optimization with the high electronic accuracy of advanced DFT methods. Compared to traditional protocols such as B3LYP/6-31G^*^, our method achieves significantly higher efficiency and better scalability to large molecules by employing fast geometric optimization with GFN2-xTB. Furthermore, by incorporating D3 dispersion correction, the method captures more accurately the long-range non-covalent interactions, such as *π*-*π* stacking and hydrogen bonding, which are often underestimated by standard DFT approaches. Additionally, compared to 6-31G^*^, the def2-SV(P) basis set provides greater accuracy, better numerical stability, and broader element coverage, while maintaining similar computational efficiency. Finally, the wave function was analyzed to extract molecular properties at multiple levels, including global, atomic, and bond-level descriptors.

### Computational scheduling and time consumption

In our scheduling system, batches of molecules are randomly assigned to different computing nodes to enable efficient large-scale computation (Figure 1). To avoid performance degradation from parallel interference, all molecular calculations are constrained to single-core execution. On each calculation node, tasks are evenly distributed across available CPU cores and executed sequentially through structure optimization, quantum property computation, wave function analysis, and feature extraction. A dynamic recycling strategy ensures full utilization of computational resources. When the overall CPU usage falls below a pre-defined threshold, completed tasks are collected and remaining molecules are redistributed for processing. This scheduling mechanism ensures near-full utilization of all CPUs across the computing cluster, thereby maximizing computational throughput. The calculations used dual Intel Skylake 6132 (14-core, 2.6 GHz) CPUs with 96 GB RAM per node, costing over 6 million CPU core hours in total.

## Results

### Overview

qcMol is a well-annotated and chemically diverse dataset designed to support drug and material discovery. This section provides a statistical overview of its key features, characterizing the composition and scope of the dataset. DFT calculations were performed at the B3LYP-D3 level with the def2-SV(P) basis set to derive the quantum chemical information. To balance computational cost, accuracy and chemical diversity, we limited the composing elements to the first four periods of the Periodic Table. The qcMol dataset comprises a total of 1,200,434 curated molecules, covering a broad spectrum of molecular sizes and weights. Each molecule is annotated with 31 quantum chemical properties, including 15 global and 16 local descriptors. The overall molecular features involve these detailed quantum chemical features as well as traditional properties computed using RDKit. Notably, 63.71% of molecules in the dataset are associated with systematic experimental results obtained through wet-lab studies, providing a valuable resource for data-driven drug discovery.

### Data characteristics

#### Data Diversity for Robust Learning

qcMol features a broad distribution in terms of elemental diversity, molecular size, and the variation of molecular scaffolds. In consideration of the salient relativistic effects in heavy elements, the database primarily focuses on non-metal elements from the first four periods of the Periodic Table (Figure 2a). The top elements in qcMol molecules include hydrogen (H), carbon (C), nitrogen (N), oxygen (O), sulfur (S) and phosphorus (P) (Figure 2b), consistent with the preference of life systems. The dataset also contains a notable number of molecules with halogen atoms, including fluorine (F), chlorine (Cl), bromine (Br) and iodine (I).

**Figure 2.**
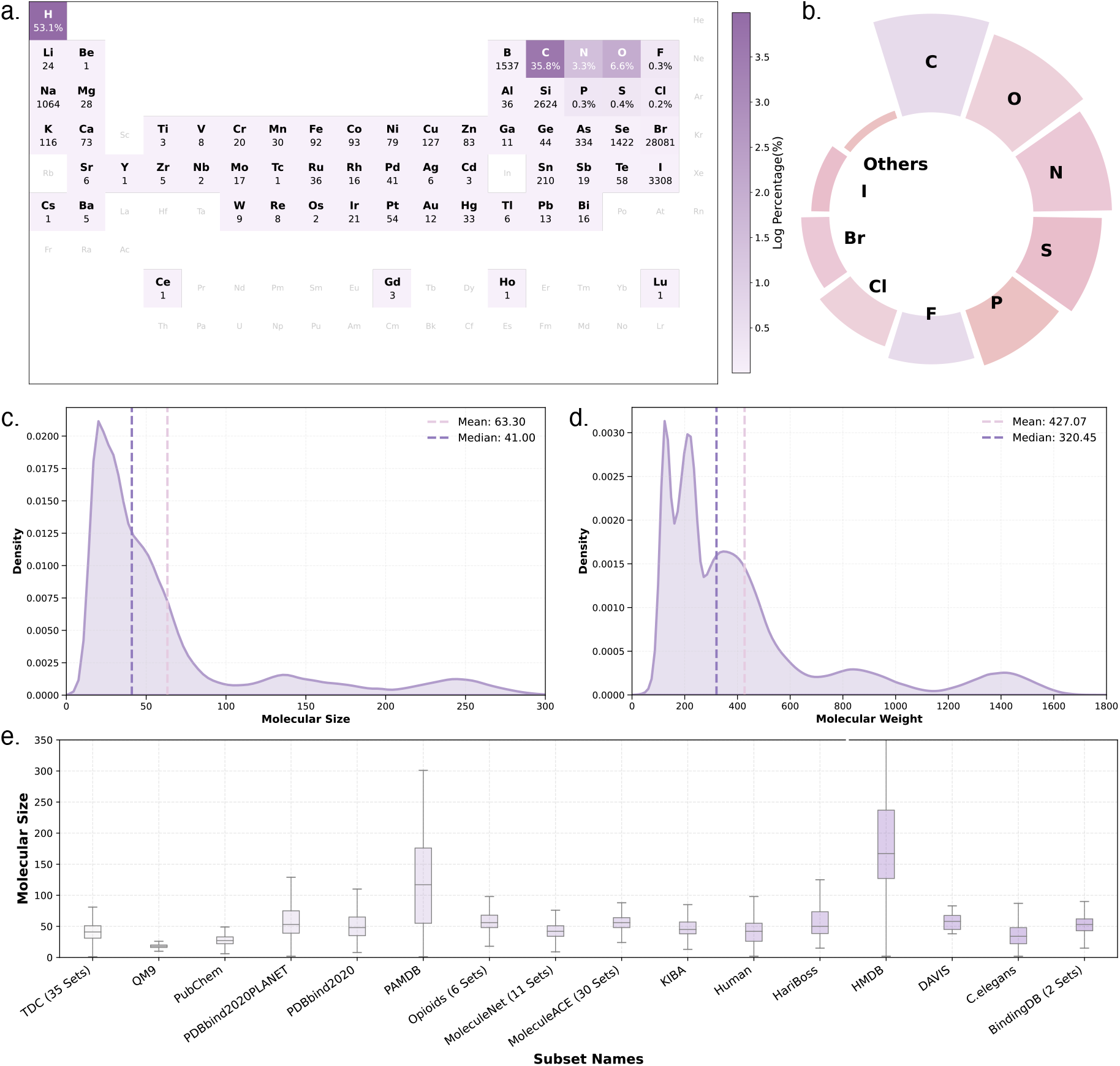
Elemental composition and molecular size distribution in qcMol. **a)** Distribution of elements involved in all existing datasets, schematically shown within the Periodic Table. Each element is colored with the degree of darkness proportional to the logarithm of its appearing percentage. The number below an element reflects the occurring count. The occurring ratio is shown instead for the top elements. **b)** Ranking of the top elements (excluding hydrogen) in qcMol. **c)** The distribution of molecular size in qcMol. **d)** The distribution of molecular weight in qcMol. **e)** Boxplot of the molecular size across different datasets, where the box limits correspond to upper and lower quartiles and the whiskers extend to 1.5 inter-quartile range.

Compared to QM9 and QuanDB, qcMol offers a wider distribution of molecular sizes (Figure 2c) and molecular weights (Figure 2d). For instance, the number of atoms per molecule in the qcMol dataset ranges from 1 to 604, covering the molecular sizes of most existing chemical datasets (Figure 2e) and thus achieving an enhanced reflection of the real-world chemical space, which is expected to be conducive to the training of robust, well-performed deep learning models.

Molecular scaffolds are widely used to split datasets for the out-of-distribution (OOD) evaluation, since molecules sharing similar scaffolds often exhibit similar properties^52^. We analyzed the scaffold diversity of qcMol and identified 247,448 unique scaffold types, with an average of 4.85 molecules per scaffold. Figure 3 provides a visualization of the most frequently occurring scaffolds, along with a quantitative analysis of the top 10 scaffold types. Notably, monocyclic scaffolds represent the dominant structural motif in the qcMol dataset, among which the most frequent ones are benzene, pyridine and cyclopropane rings. Benzene is a simple and widely occurring aromatic ring with a fully conjugated *π*-electron system, pyridine is a six-membered heteroaromatic ring commonly found in drug molecules, whereas cyclopropane is a strained three-membered ring used to restrict molecular flexibility in medicinal chemistry. These diverse monocyclic scaffolds illustrate the structural richness of the dataset and its alignment with the chemical space of drug-like compounds. Overall, the broad coverage of molecular scaffolds in qcMol makes it a potential benchmark for assessing the generalizability of molecular representation learning models.

**Figure 3.**
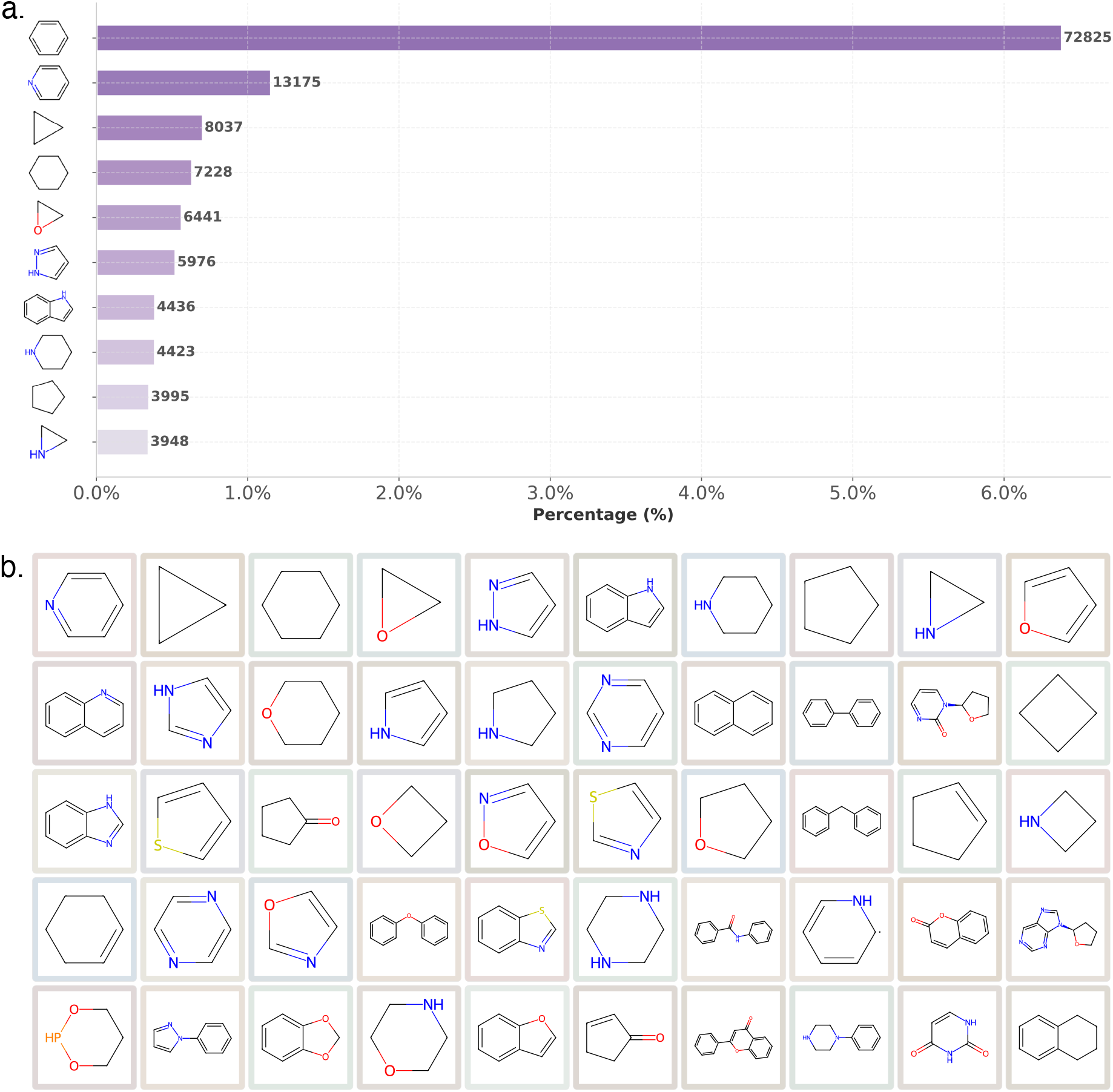
The scaffold composition of qcMol. **a)** Statistics of the top 10 scaffolds. **b)** Schematic view of the top 50 scaffolds after excluding benzene.

#### Quantum Chemical Annotation of Widely Used Molecular Datasets

Datasets like QM7, QM8 and QM9 are composed of molecules enumerated from the chemical space, many of which do not correspond to known or synthesizable compounds, weakening their applicability in real-world scenarios^32,34^. Other studies, such as the PubChemQC Project^29^, focus on calculating a large number of relatively simple molecules chosen from the PubChem database, prioritizing small-scale compounds feasible for quantum chemical calculations under the limited computational resources, which leads to the overlook of compounds with higher levels of chemical relevance or practical meaning. Unlike the above mentioned works, qcMol emphasizes the significance of the molecules being computed (Figure 4).

**Figure 4.**
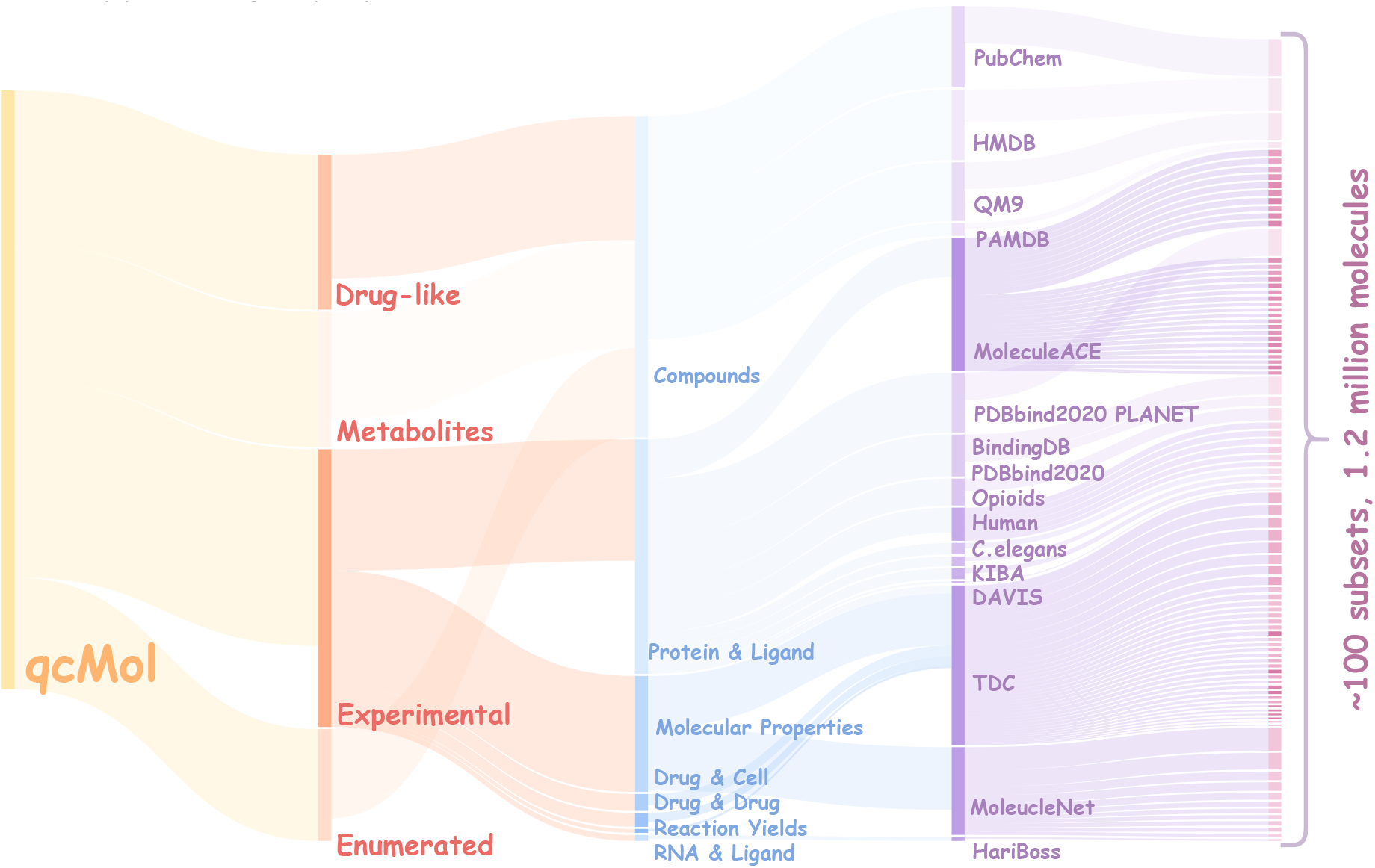
Versatile data in qcMol. Molecules in qcMol cover a broad range of compound types and are collected from a number of different datasets.

Moreover, considering the rapid data accumulation of chemical and biochemical experiments, qcMol integrates compounds from multiple datasets that capture fundamental physicochemical properties of small molecules (Supplementary Table 1). Quantum chemical annotations are especially meaningful for these molecules rather than those generated purely through computational enumeration, because molecules investigated in past experiments are often closely associated with disease processes and human physiological environment.

A key focus of qcMol is to facilitate the understanding of interactions between small molecules and biological entities, the basis on which small molecules perform specified biological functions and modulate cellular states. Accordingly, qcMol includes data covering protein-ligand interactions, RNA-ligand interactions, drug-drug interactions, and cellular responses to drugs (Supplementary Table 1).

Another important molecular category covered by qcMol falls into the class of metabolites, which are intermediate or end products of biochemical reactions within organisms and play crucial roles in maintaining cellular function, signal transduction, energy metabolism, and other physiological processes. Moreover, metabolites reflect the states of gene expression and protein activity, since they are actively involved in the regulation of internal and external cellular environments. Hence, monitoring metabolite changes allows direct detection of alterations in disease-related pathways, facilitating diagnosis and target discovery. In this consideration, qcMol incorporates metabolites in the Human Metabolism Database (HMDB) and Pseudomonas aeruginosa Metabolism Database (PAMDB) (Figure 4 and Supplementary Table 1).

In addition, this dataset also includes a set of drug-like molecules from PubChem (Supplementary Table 1). Noticeably, we have accomplished quantum chemical annotations for all molecules involved in qcMol.

In summary, qcMol currently includes five major categories of data from 95 datasets: molecular properties, intermolecular interactions, molecule-cell interactions, drug-like molecules, and metabolites. The molecules are collected from multiple different datasets, as detailed in Supplementary Table 1. By combining quantum chemical information with experimental molecular data, qcMol provides a comprehensive representation of molecules and predictive modeling for drug-like compounds.

#### Integrated Multi-Scale Features for Molecular Representation

Molecular representation learning requires that the input data be converted into formats compatible with machine learning models. Commonly used formats include strings, images and graphs, which are well-suited for natural language models^18^, computer vision models^19^ and graphbased neural networks^9^, respectively. With the growing complexity of molecular tasks, there is an increasing interest in incorporating quantum chemical information into molecular representations, enabling models to capture more detailed electronic and structural properties. Prior studies have mainly focused on global quantum chemical descriptors, such as HOMO, LUMO and the HOMO-LUMO gap, while neglecting local information at the atomic or bonding level. Although the wave function contains complete quantum information, its raw form is incompatible with most machine learning models. To address this limitation, we employed the wave function localization methods to extract structured, interpretable features at both atomic and bonding levels, enabling their effective use in downstream learning tasks. Figure 5a summarizes the main molecular features curated by qcMol.

**Figure 5.**
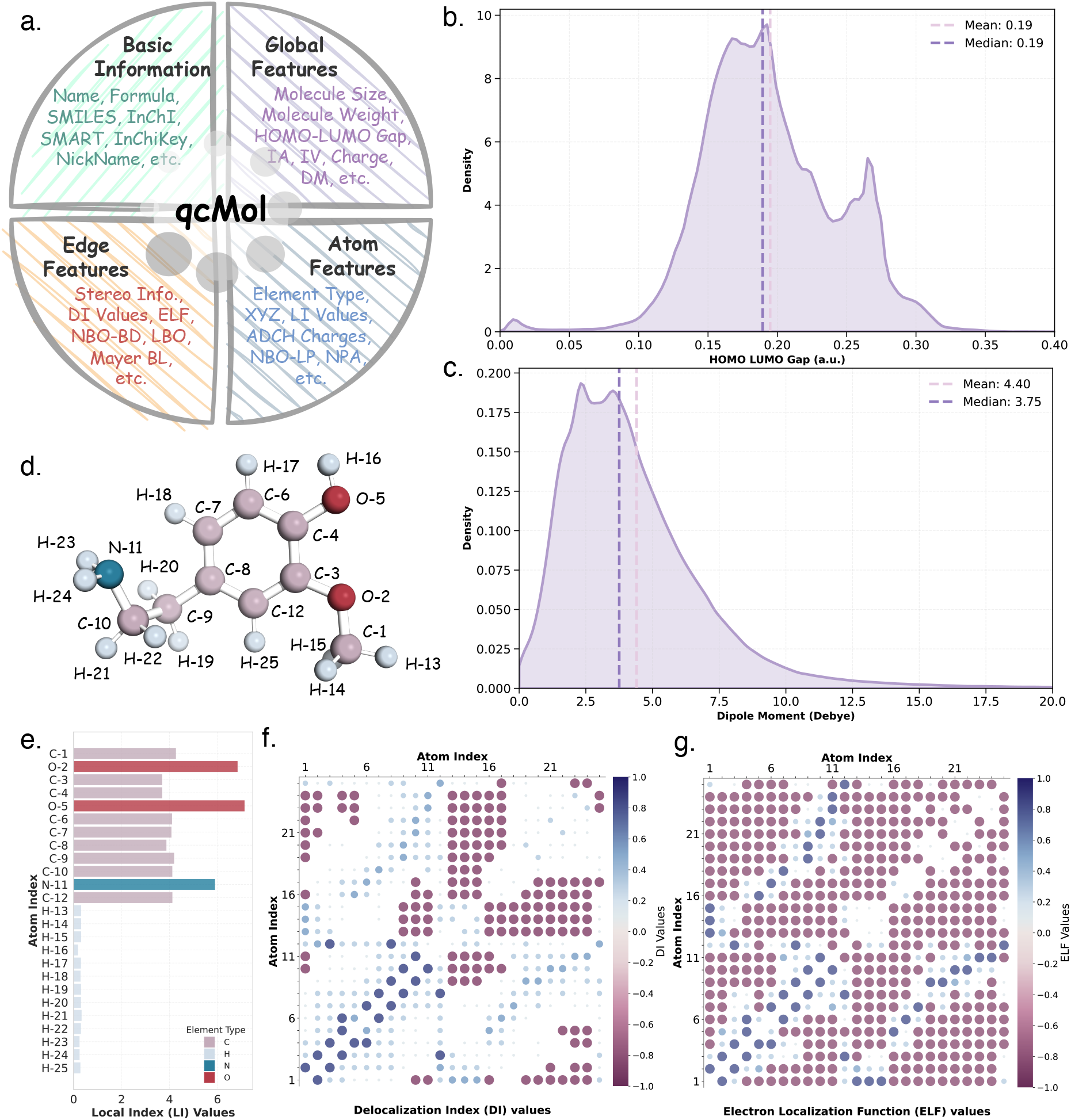
Diverse molecular features in qcMol. **a)** Overview of molecular features in qcMol. **b)** Distribution of the HOMO-LUMO gap (in atomic unit) for molecules in qcMol. **c)** Distribution of the dipole moment for molecules in qcMol. **d)** The most stable conformation of an exemplar molecule. **e)** LI representation for atoms in the exemplar molecule in panel (**d**). **f)** DI representation for bonds in the exemplar molecule in panel (**d**). **g)** ELF representation for bonds in the exemplar molecule in panel (**d**).

1. Basic information (Supplementary Table 3): This aspect includes fundamental molecular identifiers, such as qcMol ID, IUPAC name, SMILES, InChI, InChIKey, chemical formula, etc., which facilitate indexing, deduplication, and cross-database referencing.
2. Global features (Supplementary Table 4): The qcMol dataset provides 15 global features and 16 local features for the quantum chemical annotation of each molecule. The global feature includes the HOMO-LUMO gap (Figure 5b), dipole moment (Figure 5c), and other relevant properties, which primarily characterize general molecular characteristics. qcMol also includes a set of shape-related features, such as isosurface area, isosurface volume and sphericity parameters, to comprehensively capture the geometric aspects of molecular structure.
3. Local features (Supplementary Tables 5 & 6): The 16 local features consist of node-level (atom) and edge-level (bond) descriptors, organized into a molecular graph format. Exemplar node-level features are the natural atomic orbital (NAO) descriptors and local ionization (LI) values (Figure 5e), while the representative edge-level features include the delocalization index (DI) matrix (Figure 5f), electron localization function (ELF) values (Figure 5g), and Laplacian bond order (LBO).
4. Structural features (Supplementary Table 5): Previous studies have demonstrated that the three-dimensional (3D) geometric structure of a molecule plays a crucial role in determining its physicochemical properties and intermolecular interactions. In qcMol, we provide reliable geometry-optimized structures for all molecules, representing energetically stable conformations that serve as meaningful structural representations for down-stream applications (Figure 5d).

### Database Usage and Data Download

#### Web Interface and Functionality

The qcMol database is accessible through a user-friendly web interface via https://structpred.life.tsinghua.edu.cn/qcmol/. The homepage presents the core highlights of qcMol, including a clear navigation of major functionalities (Figure 6a).

**Figure 6.**
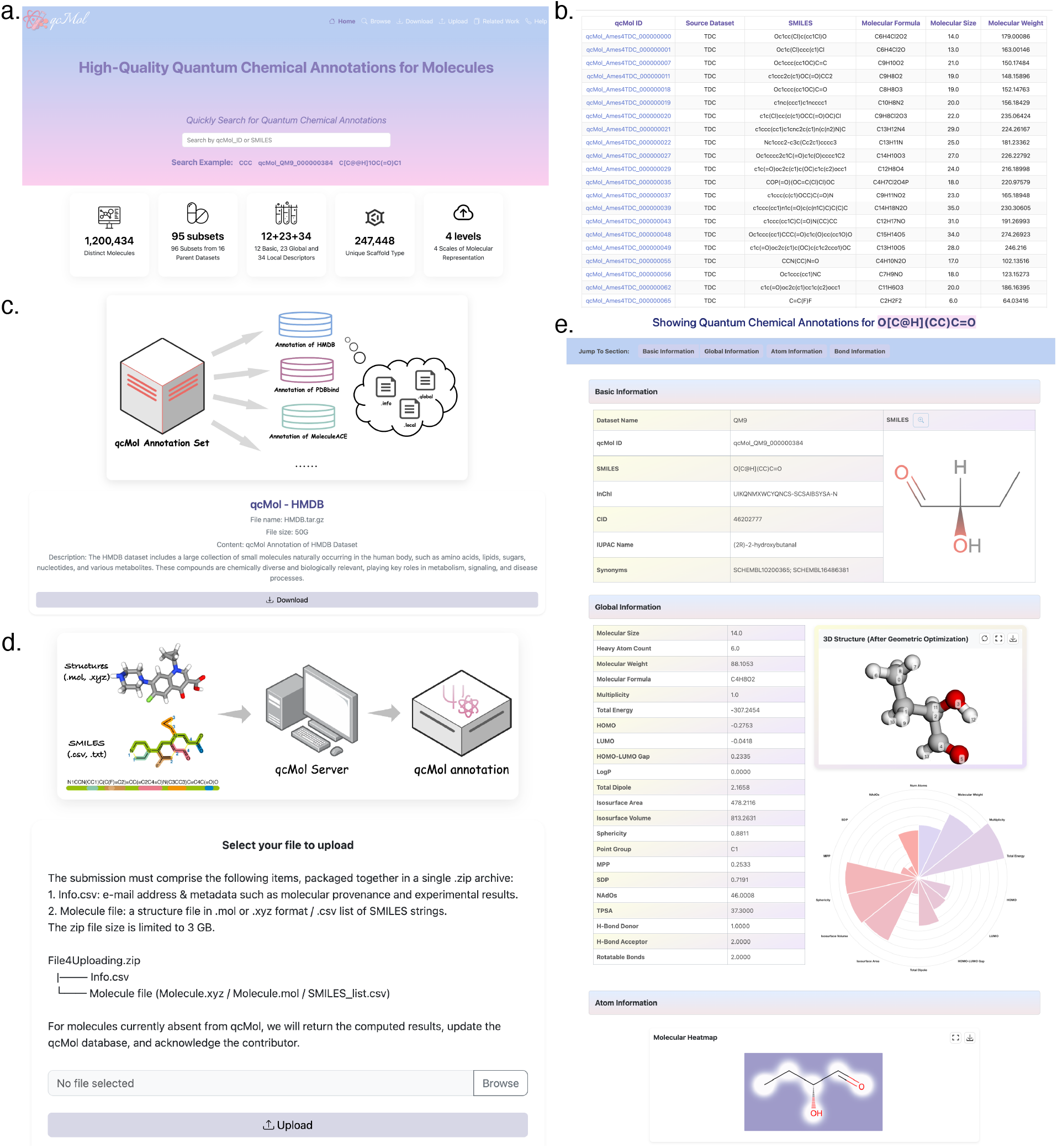
Example of the web interface for qcMol. **a)** The main interface of the web server. **b)** The browse interface of qcMol. **c)** The download interface to obtain the subset annotation. **d)** The upload interface for new molecule calculation. **e)** Exemplar page showing molecular details at the Atom Information level.

By clicking “Browse”, users can explore all molecules in qcMol and apply various filters to identify compounds that meet specific criteria (Figure 6b). The “Search” interface allows users to locate individual molecules by entering versatile identifiers including the IUPAC name, SMILES, chemical formula, InchI and other synonyms. The detailed information of each identified molecule could be visualized through further clicking (Figure 6e). The “Download” interface enables users to obtain dataset files in structured formats (Figure 6c). All raw data are publicly available for download from the qcMol website. Moreover, the dataset is further divided into multiple subsets, allowing download by specified data types. Users can also upload molecules through the “Upload” interface to request annotation of special molecules via the qcMol calculation pipeline (Figure 6d). Furthermore, the “Related Work” page will be continuously updated with recent advances and studies building on qcMol. For inquiries, please reach out via the “Help” page for more discussion.

To ensure long-term usability and expansion, qcMol will be updated every six months with newly calculated and processed molecules. Future releases will also include quantum chemical calculations for more complex molecular structures. All data are available under the Creative Commons Attribution 4.0 International License (CC BY 4.0).

#### Details of the Data Organization

We provide an example to illustrate the structure of the data after downloading. Each compound is assigned an internal qcMol ID (*e*.*g*., qcMol-Dataset-000000000). As shown in Supplementary Table 2, the molecule is stored in a dedicated folder with the following files: (1) the.info file for recording basic information; (2) the.global file to record the general properties of the molecule; and (3) the.local file to record details of atoms, bonds and structure. The following file is also available upon request for further analysis: (4) the.mol file to record the structure; (5) the.inp and.out files of the DFT calculation; (6) the.gbw and.47 files for wave function post analysis.

## Conclusion and Discussion

### Conclusion

Here, we summarize the key contributions of this work. To bridge the gap between existing quantum chemical databases and downstream applications, qcMol focuses on molecules with high chemical or biological relevance. Considering the trade-off between the limited computational resources and the vastness of chemical space, qcMol selected relevant molecules from various public datasets and performed quantum chemical calculations at the B3LYP-D3/def2-SV(P)//GFN2-xTB level. Subsequently, qcMol performed wave function post-analysis to extract structured, localized descriptors at the atomic and bonding levels.

Currently, the qcMol dataset offers quantum chemical annotations for five major categories of molecular data, including molecular properties, intermolecular interactions, molecule-cell interactions, drug-like compounds, and metabolites. This also ensures that qcMol aligns more closely with realistic chemical space by capturing diversity in molecular size, elemental composition, and scaffold structure. As a result, it promotes more robust and transferable molecular representations. By combining quantum chemical calculations with wave function analysis, qcMol provides a rich set of multi-level features, including basic information, global properties, local descriptors and geometric structures, to support diverse molecular representation learning tasks. The hierarchical organization and adaptability of these features make them especially suitable for multi-modal or hybrid modeling frameworks, which aim to comprehensively capture the structural, electronic and topological aspects of molecular systems.

Combining large-scale quantum chemical computation with feature engineering, qcMol substantially extends the scale of existing datasets and provides multi-level formats for diverse downstream tasks. This contribution is especially valuable for deepening our understanding of molecules, enhancing representation learning, and accelerating the development of novel drugs and functional materials.

### Discussion

Although qcMol offers advantages in data distribution, dataset scale and feature formats, which make it well-suited for molecular representation learning, several aspects still merit further discussion and improvement.

First, although the dataset covers a broad range of drug-like molecules, metabolites, and experimentally relevant compounds, the involved molecules only occupy a few limited local regions of the vast chemical space. The diversity in molecular size, elemental composition and scaffold structure helps improve model robustness, but rare or complex molecules remain underrepresented. Future work would focus on expanding coverage to include macrocycles, organometallics, as well as larger-sized molecules.

Second, the quantum chemical calculations were performed at the B3LYP-D3/def2-SV(P) level, which balances accuracy and efficiency well. However, more challenging tasks, like accurate reactivity prediction and excited-state property modeling, require more advanced methods such as CCSD(T) or TD-DFT.

Third, each molecule in qcMol is currently represented by a single optimized geometry. However, many molecules exhibit notable conformational flexibility, and a single structure may not adequately capture their full spatial and energetic diversity. Incorporating conformational ensembles in future dataset versions could enhance the robustness and reliability of molecular modeling, particularly for tasks sensitive to the 3D structural variability.

Lastly, how to best utilize the curated data in specific model architectures remains an open question. Systematic benchmarks and ablation studies are needed to guide model design and feature utilization strategies. Further improvements in APIs, data loaders, and integration with frameworks like PyTorch Geometric^53^ and DeepChem^54^ would enhance usability and community adoption.

## Author contribution

H. W. and H. G. proposed the concept and idea. H. W. designed the computational pipeline, collected data and performed all quantum chemical computations. Z. Z. constructed the database and the web interface. H. G. acquired funding and supervised the whole process. All authors are involved in the manuscript writing and agreed with the final manuscript.

## Acknowledgements

This work has been supported by the National Natural Science Foundation of China (#32171243), by the Ministry of Science and Technology of China (#2023YFF1204400), and by the Beijing Frontier Research Center for Biological Structure. We thank Hui Zeng for discussions and valuable suggestions.

## Supplementary Information

### Supplementary Tables

**Supplementary Table 1.**
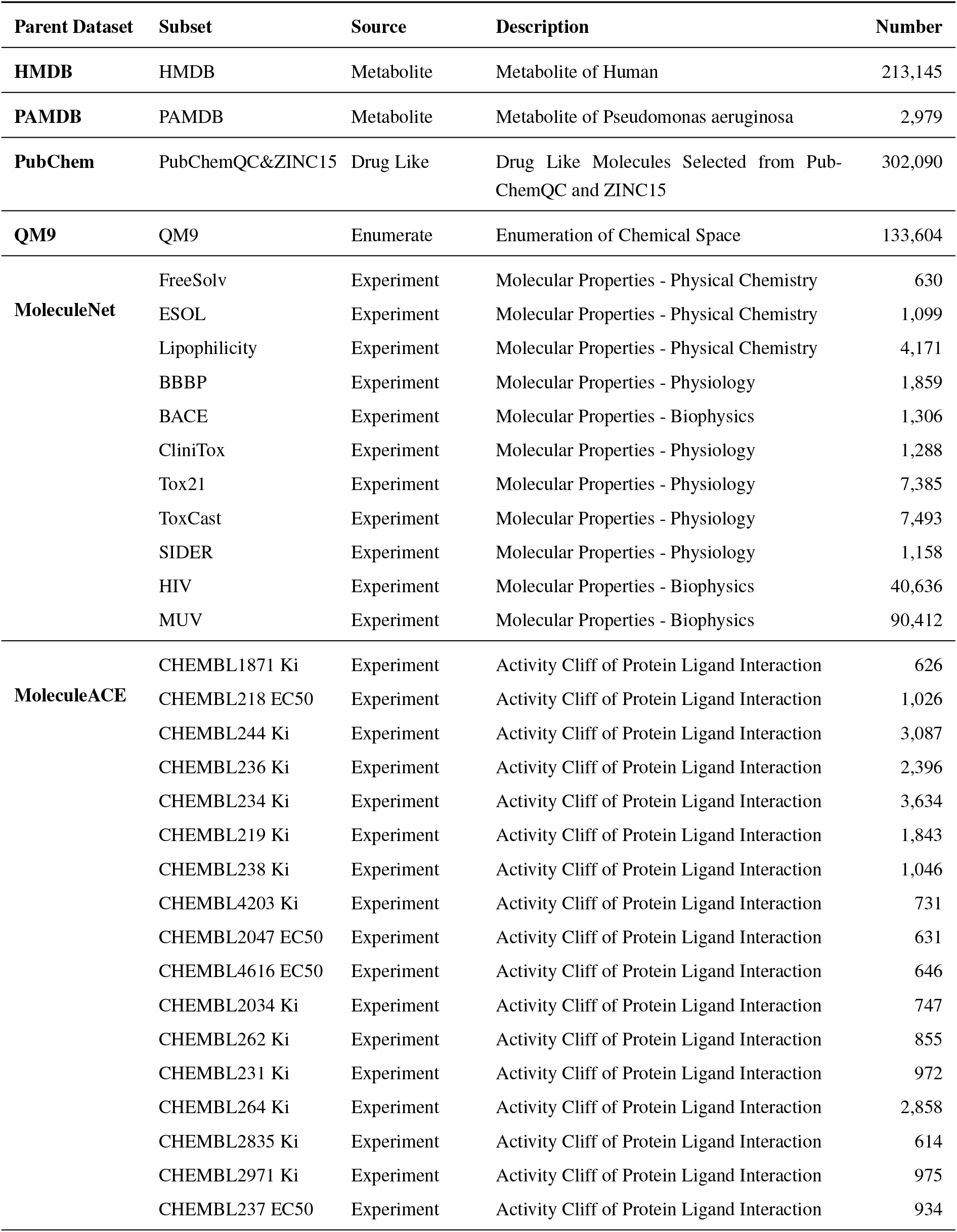
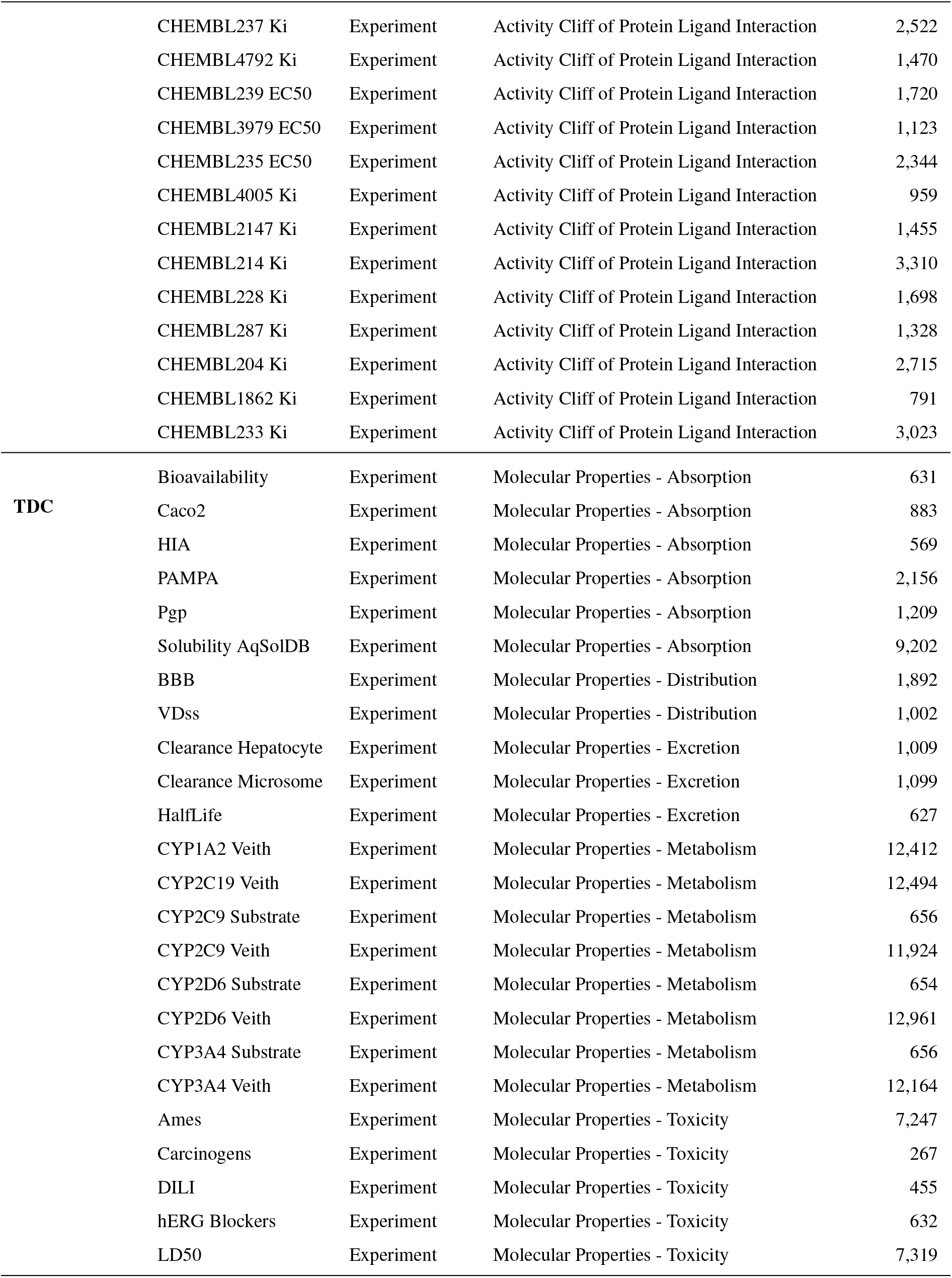
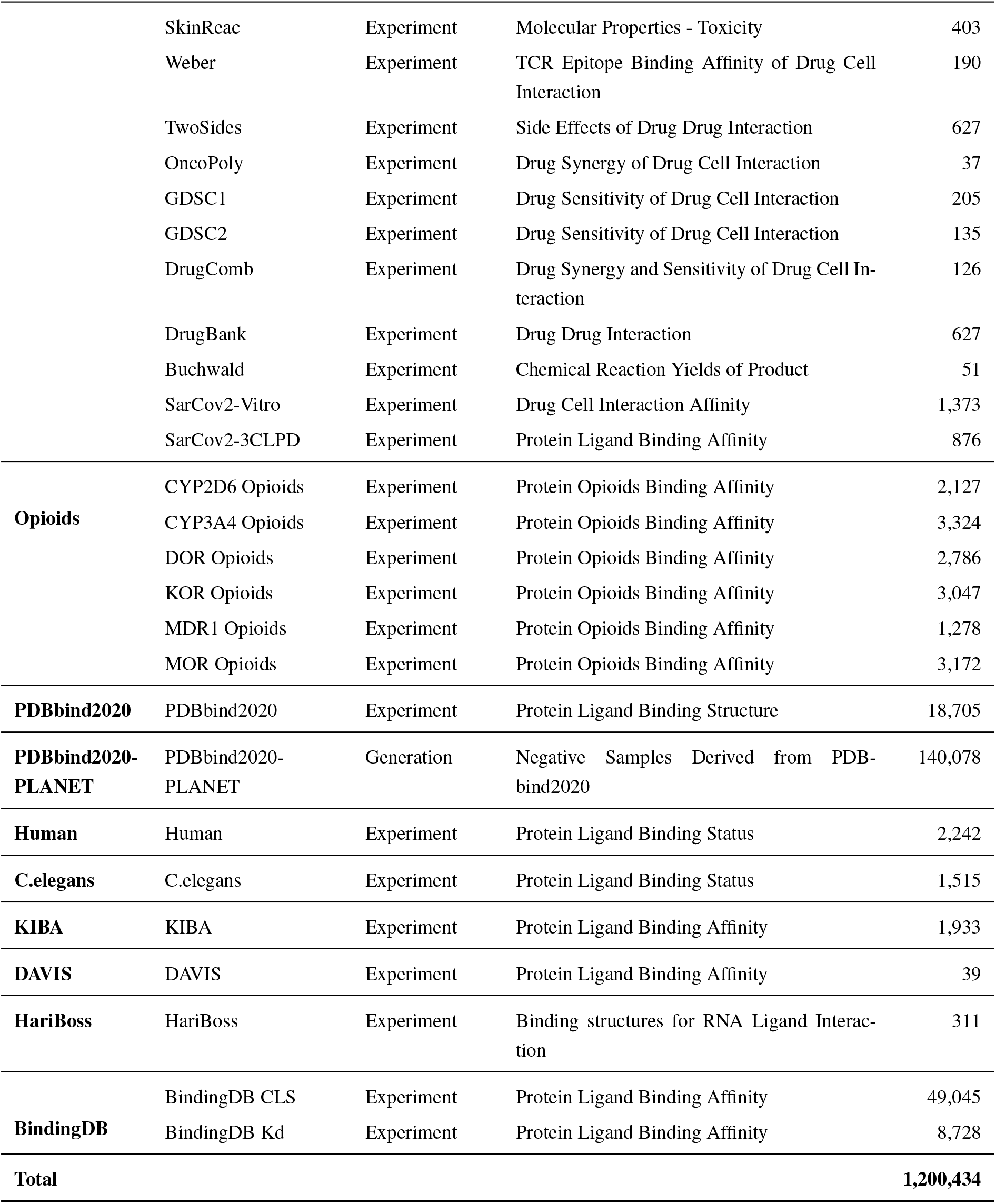
Statistics of molecular datasets.

| Parent Dataset | Subset | Source | Description | Number |
| --- | --- | --- | --- | --- |
| **HMDB** | HMDB | Metabolite | Metabolite of Human | 213,145 |
| **PAMDB** | PAMDB | Metabolite | Metabolite of Pseudomonas aeruginosa | 2,979 |
| **PubChem** | PubChemQC&ZINC15 | Drug Like | Drug Like Molecules Selected from Pub-ChemQC and ZINC15 | 302,090 |
| **QM9** | QM9 | Enumerate | Enumeration of Chemical Space | 133,604 |
| **MoleculeNet** | FreeSolv | Experiment | Molecular Properties - Physical Chemistry | 630 |
|  | ESOL | Experiment | Molecular Properties - Physical Chemistry | 1,099 |
|  | Lipophilicity | Experiment | Molecular Properties - Physical Chemistry | 4,171 |
|  | BBBP | Experiment | Molecular Properties - Physiology | 1,859 |
|  | BACE | Experiment | Molecular Properties - Biophysics | 1,306 |
|  | CliniTox | Experiment | Molecular Properties - Physiology | 1,288 |
|  | Tox21 | Experiment | Molecular Properties - Physiology | 7,385 |
|  | ToxCast | Experiment | Molecular Properties - Physiology | 7,493 |
|  | SIDER | Experiment | Molecular Properties - Physiology | 1,158 |
|  | HIV | Experiment | Molecular Properties - Biophysics | 40,636 |
|  | MUV | Experiment | Molecular Properties - Biophysics | 90,412 |
| **MoleculeACE** | CHEMBL1871 Ki | Experiment | Activity Cliff of Protein Ligand Interaction | 626 |
|  | CHEMBL218 EC50 | Experiment | Activity Cliff of Protein Ligand Interaction | 1,026 |
|  | CHEMBL244 Ki | Experiment | Activity Cliff of Protein Ligand Interaction | 3,087 |
|  | CHEMBL236 Ki | Experiment | Activity Cliff of Protein Ligand Interaction | 2,396 |
|  | CHEMBL234 Ki | Experiment | Activity Cliff of Protein Ligand Interaction | 3,634 |
|  | CHEMBL219 Ki | Experiment | Activity Cliff of Protein Ligand Interaction | 1,843 |
|  | CHEMBL238 Ki | Experiment | Activity Cliff of Protein Ligand Interaction | 1,046 |
|  | CHEMBL4203 Ki | Experiment | Activity Cliff of Protein Ligand Interaction | 731 |
|  | CHEMBL2047 EC50 | Experiment | Activity Cliff of Protein Ligand Interaction | 631 |
|  | CHEMBL4616 EC50 | Experiment | Activity Cliff of Protein Ligand Interaction | 646 |
|  | CHEMBL2034 Ki | Experiment | Activity Cliff of Protein Ligand Interaction | 747 |
|  | CHEMBL262 Ki | Experiment | Activity Cliff of Protein Ligand Interaction | 855 |
|  | CHEMBL231 Ki | Experiment | Activity Cliff of Protein Ligand Interaction | 972 |
|  | CHEMBL264 Ki | Experiment | Activity Cliff of Protein Ligand Interaction | 2,858 |
|  | CHEMBL2835 Ki | Experiment | Activity Cliff of Protein Ligand Interaction | 614 |
|  | CHEMBL2971 Ki | Experiment | Activity Cliff of Protein Ligand Interaction | 975 |
|  | CHEMBL237 EC50 | Experiment | Activity Cliff of Protein Ligand Interaction | 934 |
|  | CHEMBL237 Ki | Experiment | Activity Cliff of Protein Ligand Interaction | 2,522 |
|  | CHEMBL4792 Ki | Experiment | Activity Cliff of Protein Ligand Interaction | 1,470 |
|  | CHEMBL239 EC50 | Experiment | Activity Cliff of Protein Ligand Interaction | 1,720 |
|  | CHEMBL3979 EC50 | Experiment | Activity Cliff of Protein Ligand Interaction | 1,123 |
|  | CHEMBL235 EC50 | Experiment | Activity Cliff of Protein Ligand Interaction | 2,344 |
|  | CHEMBL4005 Ki | Experiment | Activity Cliff of Protein Ligand Interaction | 959 |
|  | CHEMBL2147 Ki | Experiment | Activity Cliff of Protein Ligand Interaction | 1,455 |
|  | CHEMBL214 Ki | Experiment | Activity Cliff of Protein Ligand Interaction | 3,310 |
|  | CHEMBL228 Ki | Experiment | Activity Cliff of Protein Ligand Interaction | 1,698 |
|  | CHEMBL287 Ki | Experiment | Activity Cliff of Protein Ligand Interaction | 1,328 |
|  | CHEMBL204 Ki | Experiment | Activity Cliff of Protein Ligand Interaction | 2,715 |
|  | CHEMBL1862 Ki | Experiment | Activity Cliff of Protein Ligand Interaction | 791 |
|  | CHEMBL233 Ki | Experiment | Activity Cliff of Protein Ligand Interaction | 3,023 |
| TDC | Bioavailability | Experiment | Molecular Properties - Absorption | 631 |
|  | Caco2 | Experiment | Molecular Properties - Absorption | 883 |
|  | HIA | Experiment | Molecular Properties - Absorption | 569 |
|  | PAMPA | Experiment | Molecular Properties - Absorption | 2,156 |
|  | Pgp | Experiment | Molecular Properties - Absorption | 1,209 |
|  | Solubility AqSolDB | Experiment | Molecular Properties - Absorption | 9,202 |
|  | BBB | Experiment | Molecular Properties - Distribution | 1,892 |
|  | VDss | Experiment | Molecular Properties - Distribution | 1,002 |
|  | Clearance Hepatocyte | Experiment | Molecular Properties - Excretion | 1,009 |
|  | Clearance Microsome | Experiment | Molecular Properties - Excretion | 1,099 |
|  | HalfLife | Experiment | Molecular Properties - Excretion | 627 |
|  | CYP1A2 Veith | Experiment | Molecular Properties - Metabolism | 12,412 |
|  | CYP2C19 Veith | Experiment | Molecular Properties - Metabolism | 12,494 |
|  | CYP2C9 Substrate | Experiment | Molecular Properties - Metabolism | 656 |
|  | CYP2C9 Veith | Experiment | Molecular Properties - Metabolism | 11,924 |
|  | CYP2D6 Substrate | Experiment | Molecular Properties - Metabolism | 654 |
|  | CYP2D6 Veith | Experiment | Molecular Properties - Metabolism | 12,961 |
|  | CYP3A4 Substrate | Experiment | Molecular Properties - Metabolism | 656 |
|  | CYP3A4 Veith | Experiment | Molecular Properties - Metabolism | 12,164 |
|  | Ames | Experiment | Molecular Properties - Toxicity | 7,247 |
|  | Carcinogens | Experiment | Molecular Properties - Toxicity | 267 |
|  | DILI | Experiment | Molecular Properties - Toxicity | 455 |
|  | hERG Blockers | Experiment | Molecular Properties - Toxicity | 632 |
|  | LD50 | Experiment | Molecular Properties - Toxicity | 7,319 |
|  | SkinReac | Experiment | Molecular Properties - Toxicity | 403 |
|  | Weber | Experiment | TCR Epitope Binding Affinity of Drug Cell Interaction | 190 |
|  | TwoSides | Experiment | Side Effects of Drug Drug Interaction | 627 |
|  | OncoPoly | Experiment | Drug Synergy of Drug Cell Interaction | 37 |
|  | GDSC1 | Experiment | Drug Sensitivity of Drug Cell Interaction | 205 |
|  | GDSC2 | Experiment | Drug Sensitivity of Drug Cell Interaction | 135 |
|  | DrugComb | Experiment | Drug Synergy and Sensitivity of Drug Cell Interaction | 126 |
|  | DrugBank | Experiment | Drug Drug Interaction | 627 |
|  | Buchwald | Experiment | Chemical Reaction Yields of Product | 51 |
|  | SarCov2-Vitro | Experiment | Drug Cell Interaction Affinity | 1,373 |
|  | SarCov2-3CLPD | Experiment | Protein Ligand Binding Affinity | 876 |
| **Opioids** | CYP2D6 Opioids | Experiment | Protein Opioids Binding Affinity | 2,127 |
|  | CYP3A4 Opioids | Experiment | Protein Opioids Binding Affinity | 3,324 |
|  | DOR Opioids | Experiment | Protein Opioids Binding Affinity | 2,786 |
|  | KOR Opioids | Experiment | Protein Opioids Binding Affinity | 3,047 |
|  | MDR1 Opioids | Experiment | Protein Opioids Binding Affinity | 1,278 |
|  | MOR Opioids | Experiment | Protein Opioids Binding Affinity | 3,172 |
| **PDBbind2020** | PDBbind2020 | Experiment | Protein Ligand Binding Structure | 18,705 |
| **PDBbind2020-PLANET** | PDBbind2020-PLANET | Generation | Negative Samples Derived from PDB-bind2020 | 140,078 |
| **Human** | Human | Experiment | Protein Ligand Binding Status | 2,242 |
| **C.elegans** | C.elegans | Experiment | Protein Ligand Binding Status | 1,515 |
| **KIBA** | KIBA | Experiment | Protein Ligand Binding Affinity | 1,933 |
| **DAVIS** | DAVIS | Experiment | Protein Ligand Binding Affinity | 39 |
| **HariBoss** | HariBoss | Experiment | Binding structures for RNA Ligand Interaction | 311 |
| **BindingDB** | BindingDB CLS | Experiment | Protein Ligand Binding Affinity | 49,045 |
|  | BindingDB Kd | Experiment | Protein Ligand Binding Affinity | 8,728 |
| **Total** |  |  |  | **1,200,434** |

**Supplementary Table 2.**
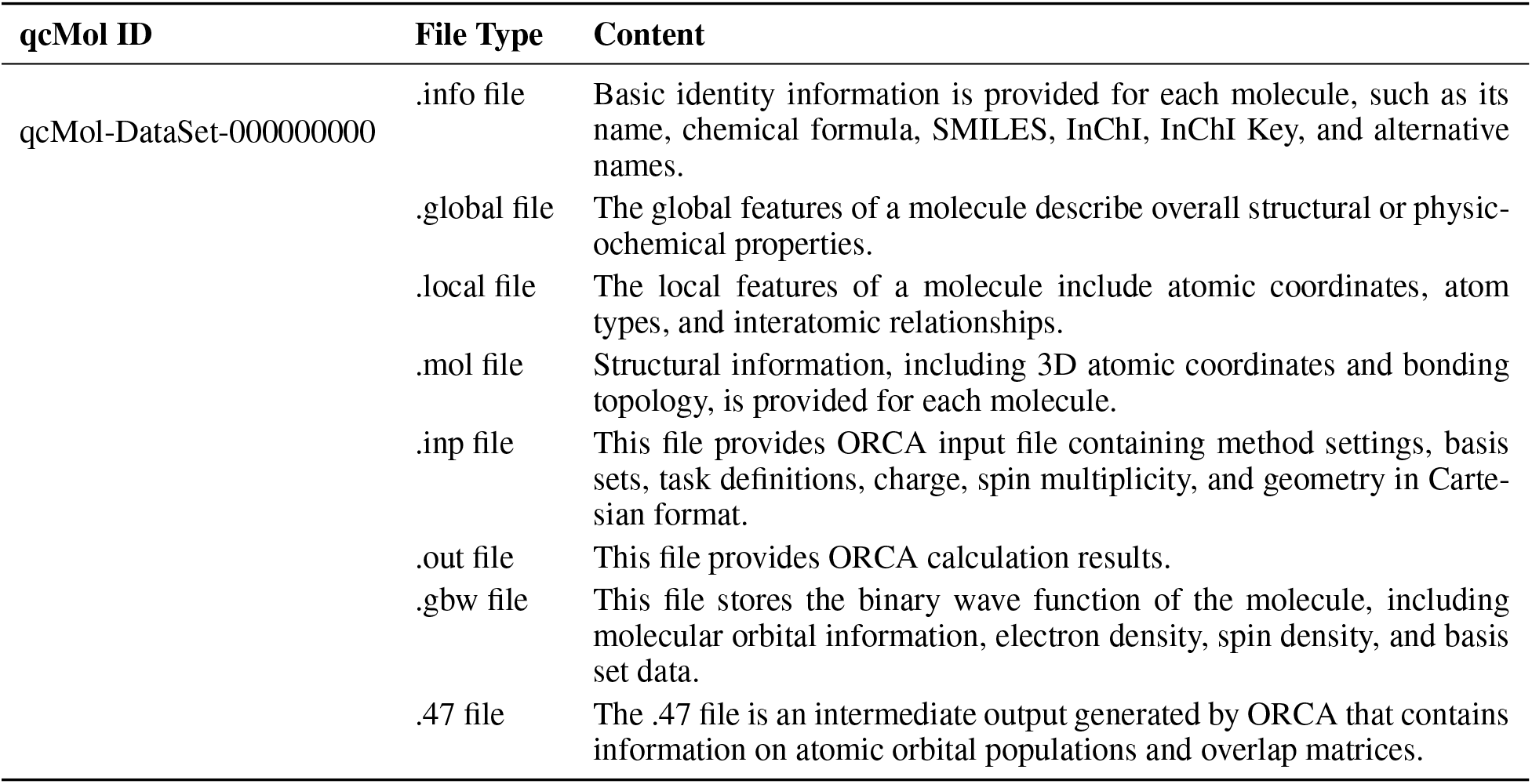
Download file structure.

| **qcMol ID** | **File Type** | **Content** |
| --- | --- | --- |
| qcMol-DataSet-000000000 | .info file | Basic identity information is provided for each molecule, such as its name, chemical formula, SMILES, InChI, InChI Key, and alternative names. |
|  | .global file | The global features of a molecule describe overall structural or physicochemical properties. |
|  | .local file | The local features of a molecule include atomic coordinates, atom types, and interatomic relationships. |
|  | .mol file | Structural information, including 3D atomic coordinates and bonding topology, is provided for each molecule. |
|  | .inp file | This file provides ORCA input file containing method settings, basis sets, task definitions, charge, spin multiplicity, and geometry in Cartesian format. |
|  | .out file | This file provides ORCA calculation results. |
|  | .gbw file | This file stores the binary wave function of the molecule, including molecular orbital information, electron density, spin density, and basis set data. |
|  | .47 file | The .47 file is an intermediate output generated by ORCA that contains information on atomic orbital populations and overlap matrices. |

**Supplementary Table 3.** Basic information for molecules.

| Type | Description |
| --- | --- |
| qcMol ID | The internal identifier of the qcMol dataset |
| Dataset Name | The source dataset for a molecule |
| CID | The internal identifier of the PubChem dataset |
| IUPAC Name | The systematic chemical name assigned according to IUPAC nomenclature rules |
| Chemical Formula | A symbolic representation that shows the types and numbers of atoms in a chemical compound |
| SMILES | The SMILES string (Simplified Molecular Input Line Entry System) from the source dataset |
| Canonical SMILES | The unique SMILES string generated in a standardized form without stereochemistry |
| Isomeric SMILES | The SMILES string that encodes stereochemistry and isotopic information |
| InChI | International Chemical Identifier |
| InChI Key | A 27-character digital identifier generated from the full IUPAC InChI, designed for efficient database indexing and web searching |
| Synonyms | An informal or common name used instead of the systematic IUPAC name |
| Version | The release version of the qcMol dataset |

**Supplementary Table 4.** Global features of molecules.

| Type | Description |
| --- | --- |
| *Basic Feature* |  |
| Molecular Size | The number of atoms in a molecule |
| Molecular Weight | The mass of a molecule |
| LogP | The logarithm of the partition coefficient between octanol and water, representing molecular hydrophobicity |
| TPSA | The topological polar surface area of a molecule, related to hydrogen bonding capacity |
| H Bond Donor | The number of hydrogen bond donor atoms in a molecule |
| H Bond Acceptor | The number of hydrogen bond acceptor atoms in a molecule |
| Rotatable Bonds | The number of rotatable single bonds in a molecule |
| Heavy Atom Count | The number of non-hydrogen atoms in a molecule |
| *Quantum Chemical Feature* |  |
| Multiplicity | The degeneracy in energy levels caused by electronic spins in a molecule |
| Point Group | Classification of molecules based on their symmetry elements |
| Total Energy | The sum of all energy terms |
| SCF Energy | Self-consistent field (SCF) energy for electrons with fixed nuclei |
| HOMO | The highest energy occupied molecular orbital |
| LUMO | The lowest energy unoccupied molecular orbital |
| HOMO-LUMO Gap | The energy difference between HOMO and LUMO |
| Mulliken | The electronic distribution within a molecule |
| Total Dipole | The overall dipole moment of a molecule |
| Isosurface Area | The surface area of a 3D contour |
| Isosurface Volume | The volume enclosed by the isosurface |
| Sphericity | The degree to which the shape of a molecule resembles a perfect sphere |
| NODOS | Nodal analysis of molecular orbitals |
| MPP | Molecular Polarizability Properties |
| SDP | Spherical Distribution Properties |

**Supplementary Table 5.** Local features of molecules (atom information)

| Type | Description |
| --- | --- |
| *Basic Feature* |  |
| Index | The atomic index in the coordinate file |
| Element Type | Chemical element symbol of the atom (*e.g.*, C, H, N, O, etc.) |
| Atomic Number | The atomic number of the corresponding element |
| Chiral Tag | The chiral information of the corresponding atom |
| Degree | The number of neighboring atoms bonded to the atom |
| Number of hydrogens | The number of hydrogen atoms directly bonded to the atom |
| Formal Charge | The formal charge assigned to each atom, based on standard valence rules |
| Radical Electrons | Number of unpaired (radical) electrons associated with the atom or molecule |
| Aromatic | Indicator of whether the atom is part of an aromatic system |
| In Ring | Indicator of whether the atom is part of a ring structure |
| Hybridization Type | The hybridization of atomic orbitals (*e.g.*, *s*, *p*, *d*, etc.) within the molecular orbital |
| *Quantum Chemical Feature* |  |
| Coordinate | Atom positions in a molecular system |
| ADCH | Atom charge distribution, reflecting the electron density distribution in the molecule |
| LI | Local ionization values, describing the atom's local ionization energy, related to molecular reactivity |
| Charge | Net atomic charge derived from the Natural Population Analysis |
| Core electrons | Number of electrons in the inner (core) shells of the atom |
| Valence electrons | Number of electrons in the valence shell available for bonding |
| Rydberg electrons | Electrons assigned to diffuse, high-energy Rydberg orbitals |
| NAO | Natural Atomic Orbital, a set of orthogonal atomic-like functions that provide an intuitive basis for interpreting molecular electronic structure. NAOs are constructed by orthonormalizing minimal basis atomic orbitals and are used to generate higher-level NBOs. |
| NBO LP | The lone-pair orbital occupied by a pair of non-bonding electrons, typically localized on a heteroatom, as defined in the Natural Bond Orbital framework. |

**Supplementary Table 6.** Local features of molecules (relationships between atoms)

| Type | Description |
| --- | --- |
| *Basic Feature* |  |
| Source Atom Index and Symbol | Chemical element symbol of the first atom in the bond |
| Target Atom Index and Symbol | Chemical element symbol of the second atom in the bond |
| Bond Type | Type of chemical bond (SINGLE, DOUBLE, TRIPLE, AROMATIC) |
| Stereo | Stereochemical configuration of the bond (STEREONONE, E, Z, etc.) |
| Conjugated | Indicator of whether the bond is part of a conjugated system |
| In Ring | Indicator of whether the bond is part of a ring structure |
| Aromatic | Indicator of whether the bond is aromatic |
| *Quantum Chemical Feature* |  |
| MBO | Mayer Bond Order matrix |
| DI | Delocalization Index matrix, quantifying electron sharing between atom pairs |
| ELF | Electron Localization Function values for each bond, measuring the spatial localization of electron pairs |
| LBO | Localized Bond Order matrix, representing the strength of bonds between atoms |
| Rigid Anchor | Rigid-atom index for FAPE loss<sup>1</sup> calculation |
| NBO BD | The Natural Bond Orbital type |
| NBO Coefficients | Series of coefficients describing the contribution of various atomic orbitals to the natural bond |

## References

1. Morgan, S., Grootendorst, P., Lexchin, J., et al. The cost of drug development: A systematic review. Health Policy 100, 4–17. ISSN: 0168-8510. https://www.sciencedirect.com/science/article/pii/S0168851010003659 (2011).

2. Rennane, S., Baker, L. & Mulcahy, A. Estimating the cost of industry investment in drug research and development: a review of methods and results. INQUIRY: The Journal of Health Care Organization, Provision, and Financing 58, 00469580211059731. https://journals.sagepub.com/doi/full/10.1177/00469580211059731 (2021).

3. Schlander, M., Hernandez-Villafuerte, K., Cheng, C.-Y., et al. How Much Does It Cost to Research and Develop a New Drug? A Systematic Review and Assessment. PharmacoEconomics 39, 1243–1269. ISSN: 1179-2027. 10.1007/s40273-021-01065-y (2021).

4. Sinha, S. K., Prasad, S. K., Islam, M. A., et al. Identification of bioactive compounds from Glycyrrhiza glabra as possible inhibitor of SARS-CoV-2 spike glycoprotein and non-structural protein-15: a pharmacoinformatics study. Journal of Biomolecular Structure and Dynamics 39, 4686–4700. 10.1080/07391102.2020.1779132 (2021).

5. Sadybekov, A. A., Sadybekov, A. V., Liu, Y., et al. Synthon-based ligand discovery in virtual libraries of over 11 billion compounds. Nature 601, 452–459. ISSN: 1476-4687. 10.1038/s41586-021-04220-9 (2022).

6. Manglik, A., Lin, H., Aryal, D. K., et al. Structure-based discovery of opioid analgesics with reduced side effects. Nature 537, 185–190. ISSN: 1476-4687. 10.1038/nature19112 (2016).

7. Lu, W., Wu, Q., Zhang, J., et al. TANKBind: Trigonometry-Aware Neural NetworKs for Drug-Protein Binding Structure Prediction in Advances in Neural Information Processing Systems 35 (Curran Associates, Inc., 2022), 7236–7249. https://proceedings.neurips.cc/paper_files/paper/2022/file/2f89a23a19d1617e7fb16d4f7a049ce2-Paper-Conference.pdf.

8. Li, Y., Hsieh, C.-Y., Lu, R., et al. An adaptive graph learning method for automated molecular interactions and properties predictions. Nature Machine Intelligence 4, 645–651. ISSN: 2522-5839. 10.1038/s42256-022-00501-8 (2022).

9. Zhou, G., Gao, Z., Ding, Q., et al. Uni-Mol: A Universal 3D Molecular Representation Learning Framework in The Eleventh International Conference on Learning Representations (2023). https://openreview.net/forum?id=6K2RM6wVqKu.

10. Feng, S., Yang, L., Huang, Y., et al. Unimap: universal smiles-graph representation learning 2023. https://arxiv.org/abs/2310.14216.

11. Friesner, R. A., Banks, J. L., Murphy, R. B., et al. Glide: A New Approach for Rapid, Accurate Docking and Scoring. 1. Method and Assessment of Docking Accuracy. Journal of Medicinal Chemistry 47, 1739–1749. ISSN: 0022-2623. 10.1021/jm0306430 (2004).

12. Halgren, T. A., Murphy, R. B., Friesner, R. A., et al. Glide: A New Approach for Rapid, Accurate Docking and Scoring. 2. Enrichment Factors in Database Screening. Journal of Medicinal Chemistry 47, 1750–1759. ISSN: 0022-2623. 10.1021/jm030644s (2004).

13. Ji, M., Gong, X., Li, X., et al. Advanced research on the antioxidant activity and mechanism of polyphenols from Hippophae species—A review. Molecules 25, 917. ISSN: 1420-3049. https://www.mdpi.com/1420-3049/25/4/917 (2020).

14. Bengio, Y., Courville, A. & Vincent, P. Representation Learning: A Review and New Perspectives. IEEE Transactions on Pattern Analysis and Machine Intelligence 35, 1798–1828 (2013).

15. David, L., Thakkar, A., Mercado, R., et al. Molecular representations in AI-driven drug discovery: a review and practical guide. Journal of Cheminformatics 12, 56. ISSN: 1758-2946. 10.1186/s13321-020-00460-5 (2020).

16. Gao, B., Qiang, B., Tan, H., et al. Drugclip: Contrastive protein-molecule representation learning for virtual screening in. 36 (2023), 44595–44614. https://proceedings.neurips.cc/paper_files/paper/2023/file/8bd31288ad8e9a31d519fdeede7ee47d-Paper-Conference.pdf.

17. Weininger, D. SMILES, a chemical language and information system. 1. Introduction to methodology and encoding rules. Journal of Chemical Information and Computer Sciences 28, 31–36. ISSN: 0095-2338. 10.1021/ci00057a005 (1988).

18. Ross, J., Belgodere, B., Chenthamarakshan, V., et al. Large-scale chemical language representations capture molecular structure and properties. Nature Machine Intelligence 4, 1256–1264. ISSN: 2522-5839. 10.1038/s42256-022-00580-7 (2022).

19. Zeng, X., Xiang, H., Yu, L., et al. Accurate prediction of molecular properties and drug targets using a self-supervised image representation learning framework. Nature Machine Intelligence 4, 1004–1016. ISSN: 2522-5839. 10.1038/s42256-022-00557-6 (2022).

20. Landrum, G., Tosco, P., Kelley, B., et al. Rdkit/Rdkit: 2025_03_2 (Q1 2025) Release 2025. http://rdkit.org/docs/Cookbook.html.

21. O’Boyle, N. M., Banck, M., James, C. A., et al. Open Babel: An open chemical toolbox. Journal of Cheminformatics 3, 33. ISSN: 1758-2946. 10.1186/1758-2946-3-33 (2011).

22. Shurrab, S. & Duwairi, R. Self-supervised learning methods and applications in medical imaging analysis: A survey. PeerJ Computer Science 8, e1045. ISSN: 2167-9843. https://europepmc.org/articles/PMC9455147 (2022).

23. Krishnan, R., Rajpurkar, P. & Topol, E. J. Self-supervised learning in medicine and healthcare. Nature Biomedical Engineering 6, 1346–1352. ISSN: 2157-846X. 10.1038/s41551-022-00914-1 (2022).

24. Kohn, W. & Sham, L. J. Self-Consistent Equations Including Exchange and Correlation Effects. Physical Review 140, A1133–A1138. https://link.aps.org/doi/10.1103/PhysRev.140.A1133 (1965).

25. Glendening, E. D., Landis, C. R. & Weinhold, F. NBO 7.0: New vistas in localized and delocalized chemical bonding theory. Journal of Computational Chemistry 40, 2234–2241. https://onlinelibrary.wiley.com/doi/abs/10.1002/jcc.25873 (2019).

26. Liu, S., Wang, H., Liu, W., et al. Pre-training Molecular Graph Representation with 3D Geometry in International Conference on Learning Representations (2022). https://openreview.net/forum?id=xQUe1pOKPam.

27. Nakata, M., Shimazaki, T., Hashimoto, M., et al. PubChemQC PM6: Data Sets of 221 Million Molecules with Optimized Molecular Geometries and Electronic Properties. Journal of Chemical Information and Modeling 60, 5891–5899. ISSN: 1549-9596. 10.1021/acs.jcim.0c00740 (2020).

28. Nakata, M. & Maeda, T. PubChemQC B3LYP/6-31G^*^//PM6 Data Set: The Electronic Structures of 86 Million Molecules Using B3LYP/6-31G^*^ Calculations. Journal of Chemical Information and Modeling 63, 5734–5754. ISSN: 1549-9596. 10.1021/acs.jcim.3c00899 (2023).

29. Nakata, M. & Shimazaki, T. PubChemQC Project: A Large-Scale First-Principles Electronic Structure Database for Data-Driven Chemistry. Journal of Chemical Information and Modeling 57, 1300–1308. ISSN: 1549-9596. 10.1021/acs.jcim.7b00083 (2017).

30. Isert, C., Atz, K., Jiménez-Luna, J., et al. QMugs, quantum mechanical properties of drug-like molecules. Scientific Data 9, 273. ISSN: 2052-4463. 10.1038/s41597-022-01390-7 (2022).

31. Yang, Z., Huang, T., Pan, L., et al. QuanDB: a quantum chemical property database towards enhancing 3D molecular representation learning. Journal of Cheminformatics 16, 48. ISSN: 1758-2946. 10.1186/s13321-024-00843-y (2024).

32. Ramakrishnan, R., Dral, P. O., Rupp, M., et al. Quantum chemistry structures and properties of 134 kilo molecules. Scientific Data 1, 140022. ISSN: 2052-4463. 10.1038/sdata.2014.22 (2014).

33. Smith, D. G. A., Altarawy, D., Burns, L. A., et al. The MolSSI QCArchive project: An open-source platform to compute, organize, and share quantum chemistry data. WIREs Computational Molecular Science 11, e1491. https://wires.onlinelibrary.wiley.com/doi/abs/10.1002/wcms.1491 (2021).

34. Blum, L. C. & Reymond, J.-L. 970 Million Druglike Small Molecules for Virtual Screening in the Chemical Universe Database GDB-13. Journal of the American Chemical Society 131, 8732–8733. ISSN: 0002-7863. 10.1021/ja902302h (2009).

35. Bannwarth, C., Ehlert, S. & Grimme, S. GFN2-xTB—An Accurate and Broadly Parametrized Self-Consistent Tight-Binding Quantum Chemical Method with Multipole Electrostatics and Density-Dependent Dispersion Contributions. Journal of Chemical Theory and Computation 15, 1652–1671. ISSN: 1549-9618. 10.1021/acs.jctc.8b01176 (2019).

36. Neese, F. Software update: The ORCA program system—Version 5.0. WIREs Computational Molecular Science 12, e1606. https://wires.onlinelibrary.wiley.com/doi/abs/10.1002/wcms.1606 (2022).

37. Neese, F., Wennmohs, F., Hansen, A., et al. Efficient, approximate and parallel Hartree–Fock and hybrid DFT calculations. A ‘chain-of-spheres’ algorithm for the Hartree–Fock exchange. Chemical Physics 356, 98–109. ISSN: 0301-0104. https://www.sciencedirect.com/science/article/pii/S0301010408005089 (2009).

38. Helmich-Paris, B., de Souza, B., Neese, F., et al. An improved chain of spheres for exchange algorithm. The Journal of Chemical Physics 155, 104109. ISSN: 0021-9606. 10.1063/5.0058766 (2021).

39. Izsák, R., Neese, F. & Klopper, W. Robust fitting techniques in the chain of spheres approximation to the Fock exchange: The role of the complementary space. The Journal of Chemical Physics 139, 094111. ISSN: 0021-9606. 10.1063/1.4819264 (2013).

40. Izsák, R. & Neese, F. An overlap fitted chain of spheres exchange method. The Journal of Chemical Physics 135, 144105. ISSN: 0021-9606. 10.1063/1.3646921 (2011).

41. Weigend, F. & Ahlrichs, R. Balanced basis sets of split valence, triple zeta valence and quadruple zeta valence quality for H to Rn: Design and assessment of accuracy. Physical Chemistry Chemical Physics 7, 3297–3305. ISSN: 1463-9076. 10.1039/B508541A (2005).

42. Lu, T. & Chen, F. Multiwfn: A multifunctional wavefunction analyzer. Journal of Computational Chemistry 33, 580–592. eprint: https://onlinelibrary.wiley.com/doi/pdf/10.1002/jcc.22885.https://onlinelibrary.wiley.com/doi/abs/10.1002/jcc.22885 (2012).

43. Wishart, D. S., Guo, A., Oler, E., et al. HMDB 5.0: the Human Metabolome Database for 2022. Nucleic Acids Research 50, D622–D631. ISSN: 0305-1048. 10.1093/nar/gkab1062 (2021).

44. Wu, Z., Ramsundar, B., Feinberg, E. N., et al. MoleculeNet: a benchmark for molecular machine learning. Chemical Science 9, 513–530. ISSN: 2041-6520. 10.1039/C7SC02664A (2018).

45. Van Tilborg, D., Alenicheva, A. & Grisoni, F. Exposing the Limitations of Molecular Machine Learning with Activity Cliffs. Journal of Chemical Information and Modeling 62, 5938–5951. ISSN: 1549-9596. 10.1021/acs.jcim.2c01073 (2022).

46. Liu, Z., Li, Y., Han, L., et al. PDB-wide collection of binding data: current status of the PDBbind database. Bioinformatics 31, 405–412. ISSN: 1367-4803. 10.1093/bioinformatics/btu626 (2014).

47. Sterling, T. & Irwin, J. J. ZINC 15 – Ligand Discovery for Everyone. Journal of Chemical Information and Modeling 55, 2324–2337. ISSN: 1549-9596. 10.1021/acs.jcim.5b00559 (2015).

48. Wang, Y., Xiao, J., Suzek, T. O., et al. PubChem: a public information system for analyzing bioactivities of small molecules. Nucleic Acids Research 37, W623–W633. ISSN: 0305-1048. 10.1093/nar/gkp456 (2009).

49. Becke, A. D. Density-functional Thermochemistry. III. The Role of Exact Exchange. The Journal of Chemical Physics 98, 5648–5652. ISSN: 0021-9606. https://pubs.aip.org/aip/jcp/article-abstract/98/7/5648/842114/Density-functional-thermochemistry-III-The-role-of?redirectedFrom=fulltext (1993).

50. Lee, C., Yang, W. & Parr, R. G. Development of the Colle-Salvetti Correlation-Energy Formula into a Functional of the Electron Density. Physical Review B 37, 785–789. https://journals.aps.org/prb/abstract/10.1103/PhysRevB.37.785 (1988).

51. Halgren, T. A. Merck molecular force field. I. Basis, form, scope, parameterization, and performance of MMFF94. Journal of Computational Chemistry 17, 490–519. ISSN: 0192-8651. https://onlinelibrary.wiley.com/doi/abs/10.1002/%28SICI%291096-987X%28199604%2917%3A5/6%3C490%3A%3AAID-JCC1%3E3.0.CO%3B2-P (1996).

52. Bemis, G. W. & Murcko, M. A. The properties of known drugs. 1. Molecular frameworks. Journal of Medicinal Chemistry 39, 2887–2893. ISSN: 0022-2623. 10.1021/jm9602928 (1996).

53. Fey, M. & Lenssen, J. E. Fast Graph Representation Learning with PyTorch Geometric in ICLR Workshop on Representation Learning on Graphs and Manifolds (2019). https://pytorch-geometric.readthedocs.io/en/latest/index.html.

54. Ramsundar, B., Eastman, P., Walters, P., et al. Deep Learning for the Life Sciences https://www.amazon.com/Deep-Learning-Life-Sciences-Microscopy/dp/1492039837 (2019).

## Supplementary References

1. Jumper, J., Evans, R., Pritzel, A., et al. Highly accurate protein structure prediction with AlphaFold. Nature 596, 583–589. 10.1038/s41586-021-03819-2 (2021).

